# A Beauty Dye Staining (ABDS) – inexpensive marker-ink based open-source alternative for commercial membrane vital dyes

**DOI:** 10.1101/2025.04.16.645980

**Authors:** Anna A. Abelit, Natalia A. Boitsova, Liudmila E. Yakovleva, Anton A. Kornev, Daniil D. Stupin

## Abstract

In this paper, we aim to present a new intravital dye, called ABDS, which can be prepared using a marker pen and is useful for eukaryotic cell research. ABDS binds to the cytoplasmic membrane of living cells and can be used for their visualization in biophysical experiments, including Z-stack tomography, cell proliferation monitoring, and adhesion studies. Important properties of ABDS are its availability, bright stable fluorescence, manufacturing simplicity, and safety for living cells *in vitro*. The paper includes a method for preparing ABDS, a data set with its characteristics, as well as examples of ABDS use in cells investigation.

## 1 Introduction

The history of science shows that important discoveries in the modern era of high technologies still can be made using mundane things and materials, which are applied in an unexpected way. Famous examples include Fleming’s invention of penicillin using poorly washed dishes; ^1^ the discovery of artificial radioactivity using Fermi’s neutron-slowing aquarium; ^2^ the invention of X-ray film medical technique by Röntgen, who observed the bones of his wife’s hands on a photographic plate; ^3^ and finally, graphene, discovered using scotch tape and a graphite pencil in 2004 by Geim and Novoselov. ^4^ Even in the advantage research of modern biology, there is a place for using everyday things. For example, nail polish is useful for fixing coverslips for microscopic and fluorescence samples, ^5–7^ a gas torch can be used to sterilize metal surfaces, ^8^ and non-fat milk is used in biochemistry protocols. ^9^ Another striking example is the use of mobile games – an inseparable part of children’s life – to assist researches related to cancer (Cancer Research UK games, Genima) ^10,11^ and protein folding (FoldIt). ^12^

Frequently, scientists’ motivation to improvise with methods and materials at hand is caused by a desire to increase research cost-efficiency, speed up study progress, or due to the absence of available on the market required chemicals or apparatus. A bright example of such research optimization in biology was done in 2013 by Yuta Takase *et al*. who have used highlighter ink to visualize blood vessels in avian and mouse embryos using fluorescent microscopy. ^13^

In 2021, the method of visualizing blood vessels using yellow highlighter ink was used in a study by Luke Bollinger and Renee Dickie, who studied axolotl regeneration. ^14^ They also improved this imaging method by making it possible to image vessels using a fluorescent adapter system with a stereomicroscope and a phone camera. Finally, the visualizing vessels using highlighter ink was also used in a 2023 study by Mijithra Ganesan *et al*., in their study of regeneration in earthworms. ^15^ This study did not use fluorescence microscopy, but did visualize vessels using highlighter ink, allowing them to be observed with a light microscope.

In our study, following this line, we expand the area of such a “lab-scale” concept by improving the idea of the writing implements utilization in biology, and propose a cell membrane vital dye, which is prepared from commercially available permanent marker ink. Due to the outstanding property of cell membrane visualization, we named our dye ABDS as the abbreviation of “A Beautiful Dye for Staining”. Contrary to works ^13–15^ our staining technique allows to investigate micro-objects such as single living cells in real time mode with submicron resolution.

We present (i) the preparation protocol of ABDS including a 3D printed stencil to control the concentration of ABDS, (ii) the study of the optical properties of ABDS including absorption, fluorescence and Raman spectra, (iii) the demonstration of the applications of ABDS in living cell research, including the comparison of ABDS with a low-cost cell membrane dye DiBAC (ThermoFisher Scientific, USA). Using the obtained data, we drawn conclusions about its composition and, furthermore, we found the effect of gradual increase in brightness of ABDS during long-term optical pumping, which can be explained by quantum mechanics. Compared to most commercially available dyes, ABDS is cheaper, so it will be useful for biological laboratories that often face the need for a simple and inexpensive method to visualize live cells, ^16^ as well as for do-it-yourself (DIY) biological devices such as smartphone-based microscopes. ^17^ We believe that ABDS will optimize the research activities of most biological laboratories, which will stimulate progress in biology, cytology, and medicine.

The article is organized as follows. First, we present an ABDS preparation protocol and introduce various characteristics of ABDS such as its absorption, emission, and Raman spectra. Then, we will consider the interaction of the ABDS dye with two eukaryotic cell lines, such as HeLa and RIN m5F, and demonstrate unique ABDS features. Next, we will compare the properties of ABDS with those of the other intravital dyes – DiBAC and Deep Red Cell Mask (ThermoFisher Scientific, USA). After that, we will discuss the application of these features of ABDS in biology and medicine with special attention payed to cell proliferation research. Finally, we will provide examples of the use of ABDS in impedance spectroscopy ^18^ to study phenomena that occur in cell populations and single cells.

## 2 Materials and Methods

### 2.1 ABDS preparation

To prepare ABDS the 96% ethanol, a glass slide, PBS, and a permanent red marker (like red Edding E-140S, Fig. 1) are required. The detailed preparation protocol is as follows (Fig. 2):

**Figure 1:**
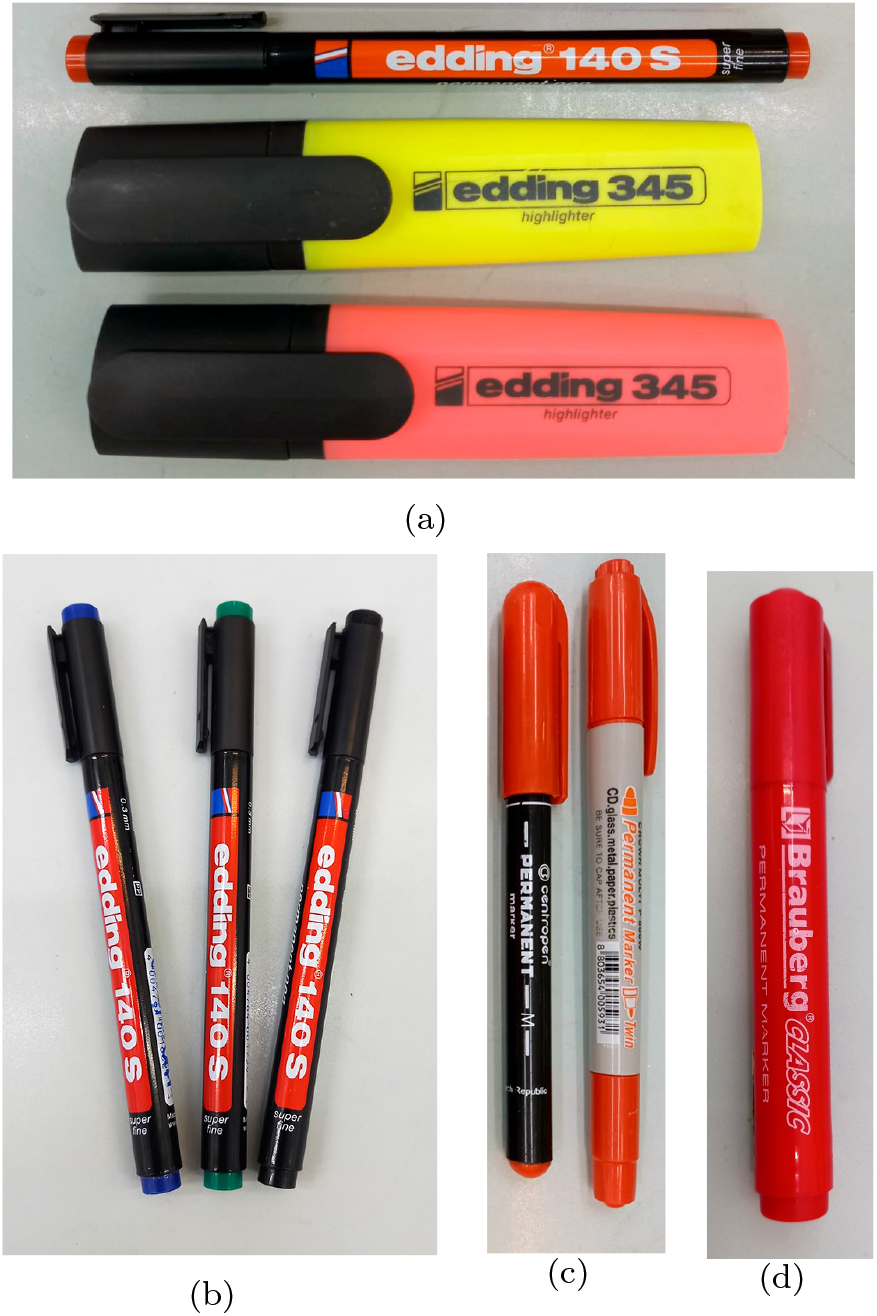
Markers types that we have tested.

**Figure 2:**
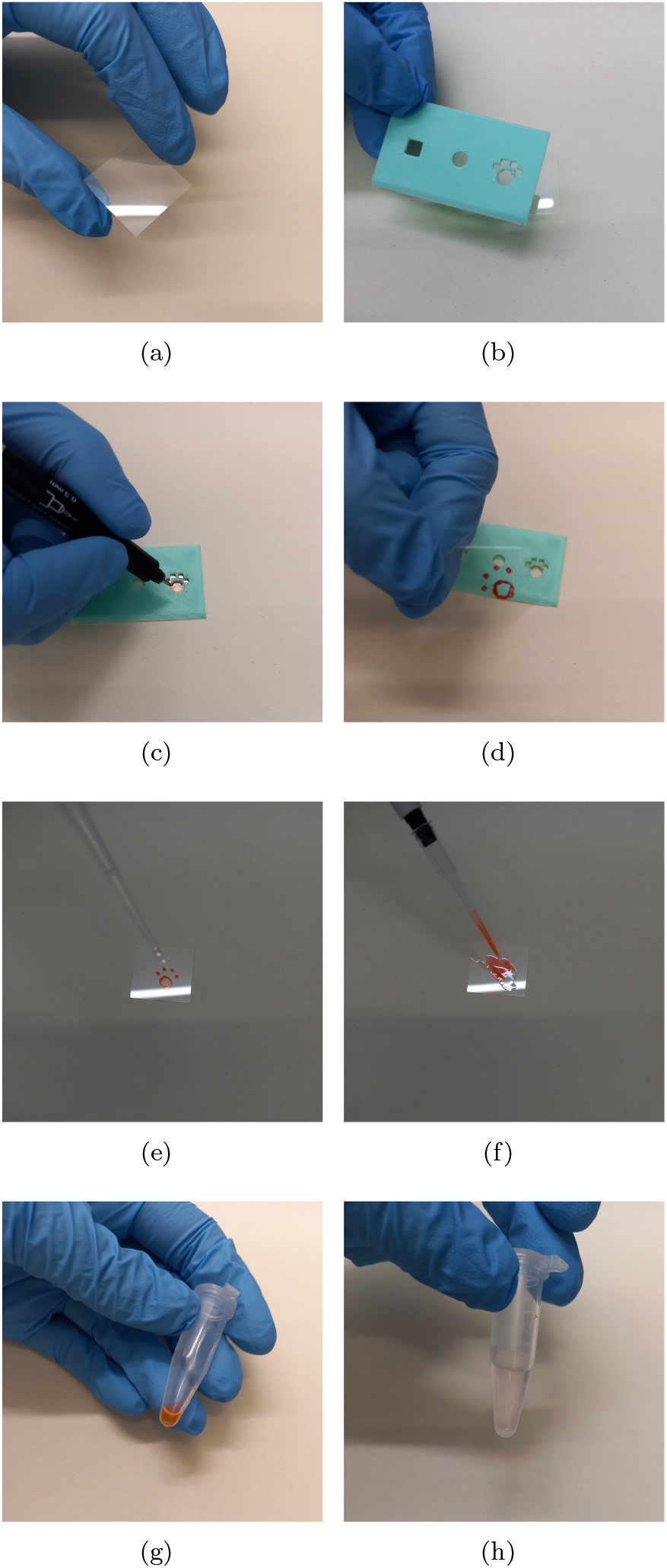
ABDS Preparation Protocol: (a) – glass slide, (b) – 3D printable spencil, (c) and (d) – drawing picture with red permanent marker, (e) and (f) – dissolving the picture with 50 *µ*l of 96% ethanol, (g) and (h) – dilution 5 *µ*l of the resulting ethanol solution in 1 ml of PBS.

1. In order to unify the amount of ink used for the dye, draw an open square of size 4 *×* 4 mm^2^ on a glass slide [Fig. 2(a-d)] or any other pattern. The length of the line should be between 20 and 50 mm. We recommend using special 3D-printed stencil, stl-file for which can be obtained by request or found in the this article source code.
2. Next, dissolve the depicted pattern with 50 *µ*l of 96% ethanol [Fig. 2(e-f)].
3. After this, it is necessary to dilute 5 *µ*l of the resulting ethanol solution in 1 ml of PBS [Fig. 2(g-h)].

Preliminary we have tested different types of markers (Fig. 1) including black, green, and blue permanent Edding E140S, red whiteboard marker Crown Multi WB-505 (Crown, Korea), and yellow and pink highlighters Edding 345 (Edding International GmbH, Japan). We found that bright fluorescence of cells was observed only when using red permanent markers. We tested permanent red markers from various manufacturers and the results were reproducible with all of them, suggesting that the red permanent marker ink uses the same active ingredient. The ABDS-like dye, prepared from pink and yellow highlighter pens, demonstrate fluorescence effect significantly lower than red permanent markers, thus we choose the red Edding E-140S pen in this article. Therefore for ABDS preparation it is very important to use red and permanent markers, because inks with other colors and inks for whiteboard markers do not provide a fluorescence effect. This can be the result of the absence of the Rhodamine 6G component in them, which we assume provides a staining effect in the cells (see Sec. 3.2). In this study, we have mostly used red Edding E-140S (edding International GmbH, Japan) marker with linewidth of 0.3 mm [Fig. 1(a), upper marker].

The ABDS dye is resistant to aggregation. However, for ABDS storage we recommend using glass tubes or using plastic tubes, but for no longer than 1 hour, since the dye tends to be absorbed into plastic.

### 2.2 3D printable stencil

Since the initial concentration of the marker ink is typically unknown, we had to create a special stencil to control the concentration of ABDS, which has a square hole 5 *×* 5 mm^2^, a circle hole with 5 mm diameter and a cat paw hole, which we used for ABDS preparing (Fig. 3). Also, the stencil has a rule cut for making an arbitrary length market ink line. The stencil is used for creating ABDS with reproducible concentration, *e*.*i* it allows us to control marker ink amount by marker line length (for square hole the line will be have approximately 20 mm length, for circle hole it will be 5*π* mm, arbitrary length can labeled on the rule cut). The stencil was developed in Blender 4.3 software and printed from PLA (Guangzhou Zhiwen Technology, China) on the Flying Bear S1 3D printer (Maylerescape, UK).

**Figure 3:**
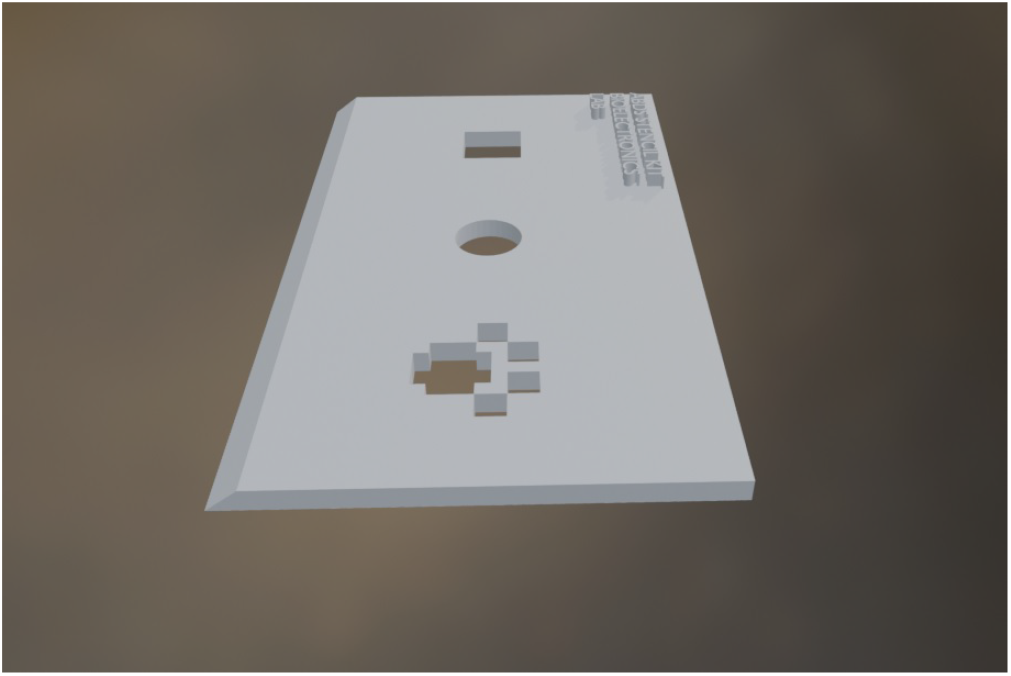
3D printable stencil. Stl-file can be found in the article source code, obtained by request or download from link. ^19^

### 2.3 Optical spectroscopy study

In order to estimate the ABDS composition, we have measured its absorption, emission, and Raman spectra. The absorption was measured on the Evolution 300 UV-VIS spectrophotometer (ThermoFisher Scientific, USA) in the 400-800 nm range. The emission spectra were collected using a Chirascan spectrophotometer (Applied Photophysics, UK) using an excitation wavelength of 525 nm. The Raman spectra were obtained with the LabRam HR800 setup (Horiba, France) using IR laser 784.19 nm.

### 2.4 Cell cultures

In this study, we have used HeLa and RIN m5F cells, obtained from the Bank of cell cultures of the Institute of Cytology of the Russian Academy of Sciences. For cell cultivation, we used the DMEM medium (Biolot, Russia) for HeLa and the RPMI medium (Biolot, Russia) for RIN m5F. For both cell lines, 10% fetal bovine serum (HyClone, USA) and antibiotic gentamicin (Biolot, Russia) were added to cells medium before incubation, which was performed at 37^*°*^C and 5% CO_2_.

### 2.5 Intravital staining of the cells

Intravital cells staining with ABDS was carried out as follows. First, it is necessary to wash the cells from the culture medium with PBS at least twice. Then a PBS-solution of ABDS needs to be added to the cells (the preparation protocol is presented in Sec. 2.1) after which cells should be incubated with the dye solution for 10 minutes at 37°C. Finally, the cells should be again washed with PBS twice. For cells immediate optical research, we recommend using optically transparent PBS as cells medium. Also, it is possible to incubate stained cells in culturing medium if it is planned to study them over several days. In this study, we have used analogical protocol for cells DiBAC staining.

### 2.6 Microscopy and third-party cell membrane dyes

For ABDS staining examination we have used Leica 4000 DM B florescence microscope (Leica, Germany, A, I3, and N21 cubes) and Zeiss Observer.Z1 confocal microscope (Zeiss, Germany) equipped with pumping lasers source with wavelength 532/561 nm and 488 nm (Zeiss Lasermodul TIRF 230V, Germany). The Zeiss photographs were processed using AxioVision (SE64 Rel. 4.9), since the data obtained from the Zeiss microscope contains a larger than 256 number of tones (4096), which does not allow them to be processed using Adobe Photoshop (26.2.0), like photographs from a Leica microscope. The images were collected as 2D photographs and as 3D Z-stack tomography. For high-resolution confocal pictures, immersion oil was used. As a sample for comparison, we have used commercial low-cost dye DiBAC and moderate cost dye Deep Red Cell Mask (ThermoFisher Scientific, USA). The photographs are presented in monochrome or pseudo-color modes.

### 2.7 Viability test

The viability test of the ABDS was performed using the cell BD FACSCanto Flow Cytometer (BD Bioscience, USA) and DAPI (dead cells nuclei visualization, ThermoFisher Scientific, USA) dye for dead cell detection. To monitor cells viability during the microscope study, we have also used the DAPI dye. All microscope photographs were made in the optically transparent non-fluorescence phosphate buffered saline (PBS, RosMed-Bio, Russia).

## 3 ABDS properties and composition

### 3.1 ABDS florescence study using microscopes

To study ABDS cells staining feature, we have provided a wide range of experiments with fluorescence microscopy. At first, we have exanimate an existence of the ABDS fluorescence in HeLa cells with a Leica 4000 DM B florescence microscope (Leica, Germany). The results obtained for cubes I3, N21 and A, as well as a reference bright-field image of the cells are shown in Fig. 4, which clearly shows that ABDS successfully stains the cell membrane and allows to obtain high-quality cells images. Photographs of cells in Fig. 4 were processed with the highest image clarity using the “curves” tool in Adobe Photoshop 26.2.0 (shape of the curves for each of the photographs is shown bottom panels in Fig. 4). It can be seen that the ABDS photograph obtained with cube N21 (green light pumping, red light emission) has the best image quality, the photograph obtained with cube I3 (blue light pumping, green light emission) has less contrast, and cube A with UV pumping has very weak ABDS signal. These results show that the green light range is preferable for ABDS pumping. It also follows from Fig. 4 that ABDS is compatible with UV-pumped blue dyes such as DAPI.

**Figure 4:**
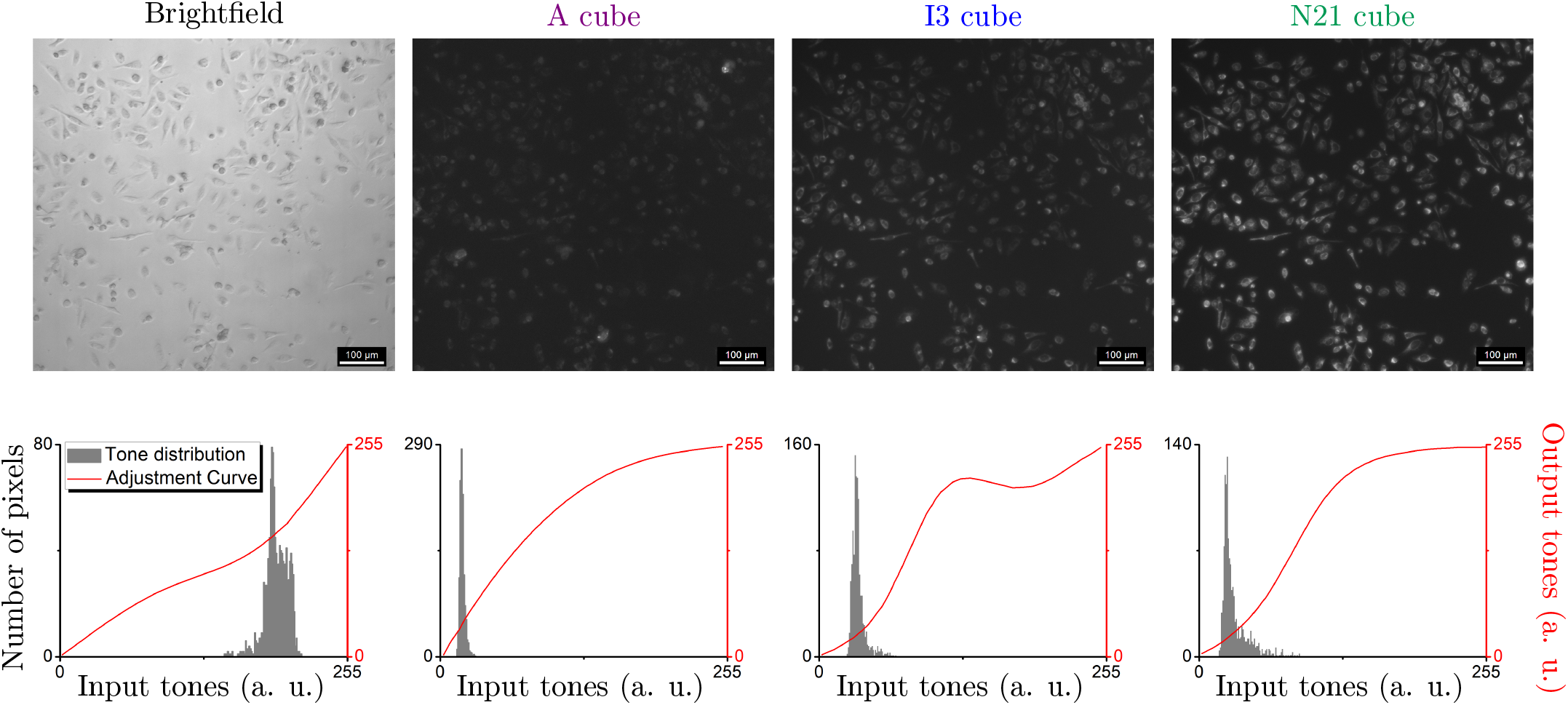
ABDS fluorescence in HeLa cells with Leica 4000 DM B florescence microscope. With this microscope fluorescence were studied using I3, N21, and A cubes. The photographs’ tones were processed by Adobe Photoshop.

The more striking result of the ABDS HeLa staing is shown in Fig. 5, where obtained with Zeiss microscope confocal images of the HeLa cells treated with ABDS are presented. These photographs demonstrate the submicron resolution of the ABDS dye, which allows visualization of not only membrane topology but also cytoskeletal structures. Similar to the Leica data, the Zeiss raw images were adjusted with AxioVision software for better clarity. From the adjusted images in Fig. 5 it can be concluded that pumping in the green and blue light lasers are suitable for ABDS excitation. In addition, we also observed no ABDS signal with usage of red laser pumping 639 nm, which makes ABDS compatible with IR fluorescence dyes like Deep Red Cell Mask.

**Figure 5:**
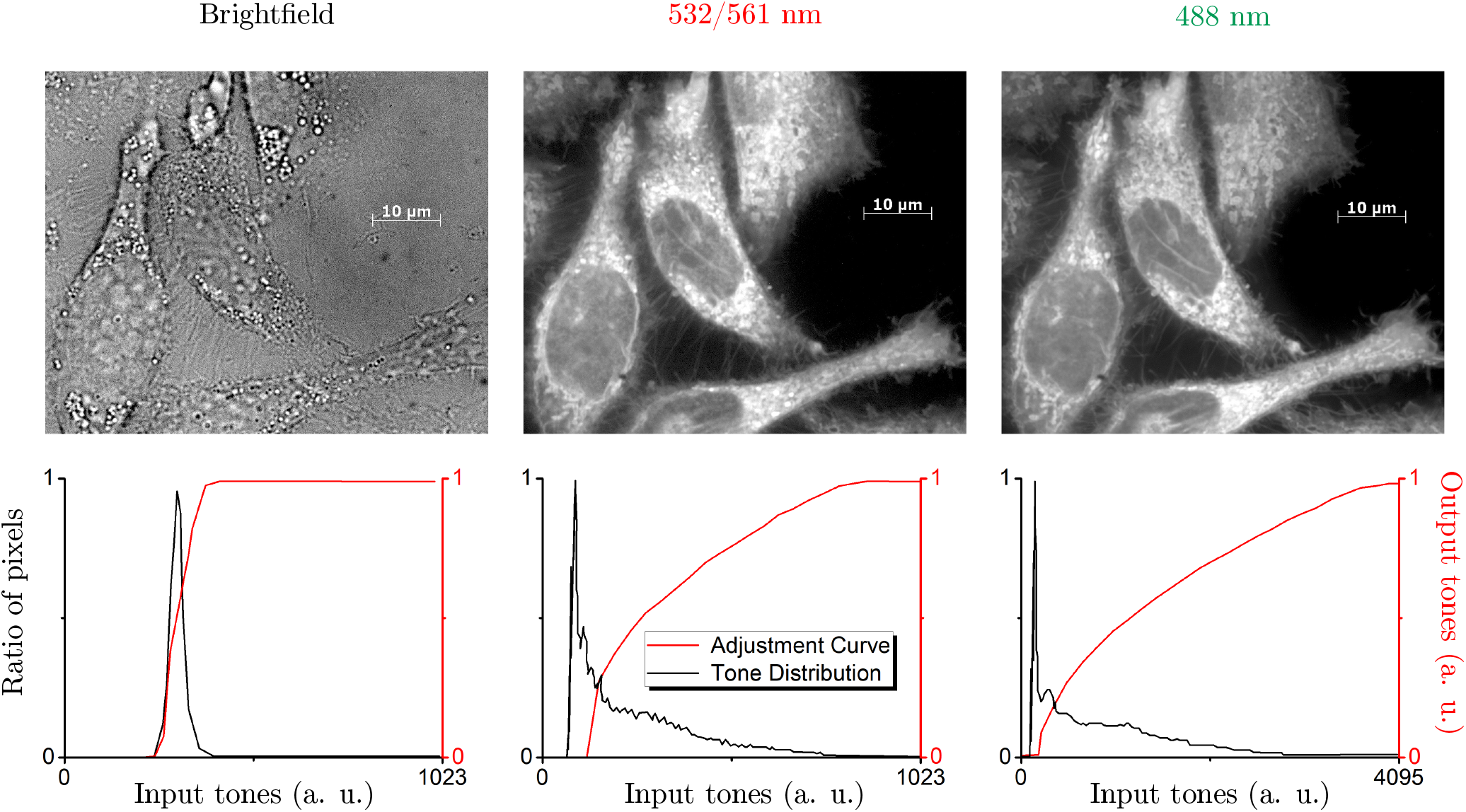
ABDS fluorescence in HeLa cells with Zeiss Observer.Z1 confocal microscope. With this microscope fluorescence were studied using brightfield, and in fluorescence mode with lasers 532/561 nm and 488 nm. The photographs were processed by AxioVision (tuning curves are presented at bottom panels). The obtained data indicates that it is possible to study cells’ membrane structure with submicron resolution using ABDS. The difference in the tone range in histograms for lasers can be due to different lasing light intensity, which is typically unknown and very difficult to measure.

### 3.2 ABDS absorption, emission, and Raman spectra

Since the exact composition of the Edding 140-S permanent red marker ink is unknown and most likely is a trade secret, we attempted to reverse engineer the fluorescent substance in the ink. In order to determine which substance fluoresces in the composition of ABDS, we examined it using absorption, emission, and Raman spectroscopy. The results are shown in Fig. 6 and 7. One can see, ABDS has two absorption peaks at 528 nm and 675 nm, and it’s Raman spectra demonstrate distinct peak signature. After analyzing ABDS absorption spectra (Fig. 6) and it’s fluorescence (Fig. 4 and Fig. 5), we conclude that ABDS has a pumping level near 528 nm. Using this fact, we have measured the emission spectra pumping ABDS near this level. As a result, we observed the 50-nm width emission peak at 552 nm, *e*.*i*. in the green-red color range, which we can see using our eyes on the microscope (Fig. 4 and Fig. 5).

**Figure 6:**
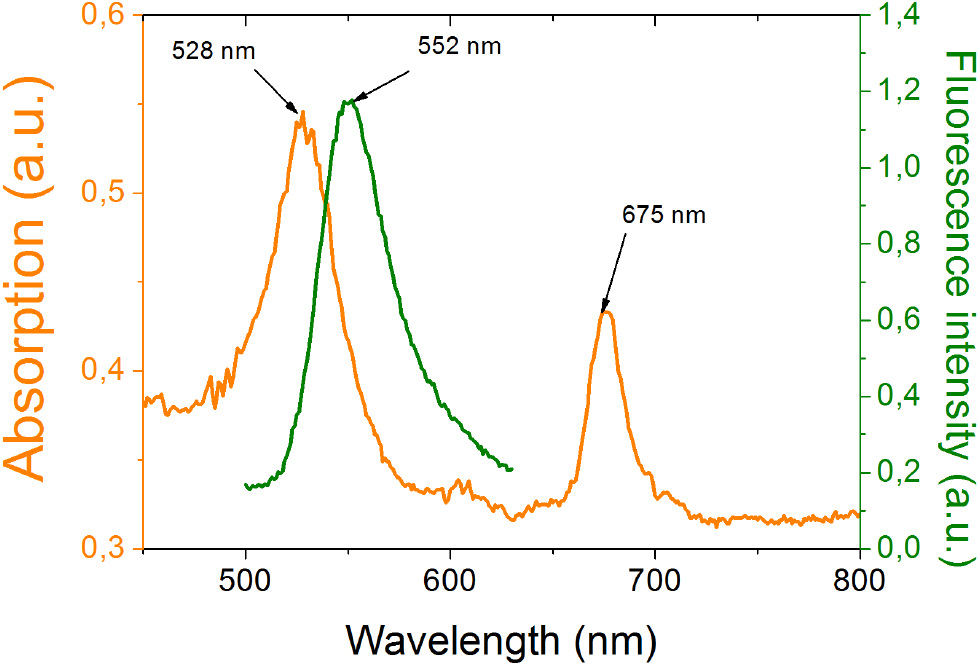
Absorption (orange) and emission (green) ABDS spectra.

**Figure 7:**
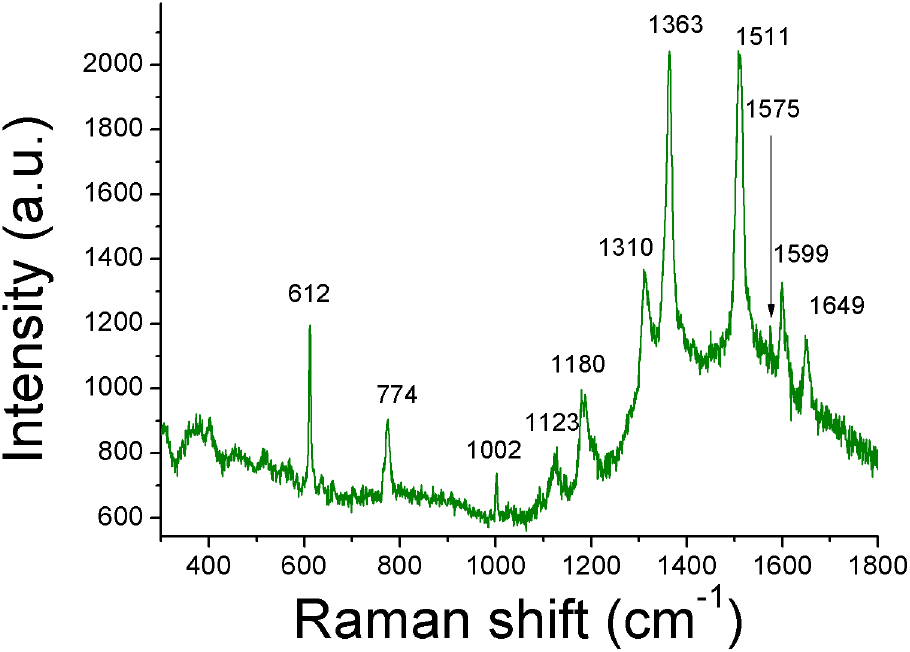
Raman spectrum of ABDS. The peaks signature is close to Rhodamine 6G. ^24^

Since fluorescent staining could be carried out not only with the help of the marker Edding 140-S, but also with red fluorescent markers from other companies, we concluded that the substance used as a dye is widely used. A search in various literary sources led to the assumption that the coloring pigment of red alcohol ink or one of its components is the substance Rhodamine 6G. ^20,21^ At the same time, the absorption, ^22^ emission, ^23^ and Raman spectra are known for Rhodamine 6G, ^24^ and coincide with the ABDS spectra. Thus, we can assume that Rhodamine 6G is present in the composition of the ABDS dye. Possible additional peaks in the Raman and absorption spectra of the ABDS may mean that the ink of the permanent red marker also contains other substances, which we, however, had not identified yet.

## 4 ABDS features

ABDS has several undeniable advantages, including high cell survival after staining, stable fluorescence, and low-cost. This section demonstrates these features in detail.

### 4.1 Cytotoxicity test

In order to determine the cytotoxicity of ABDS, a flow cytometric study was conducted. The dye-containing samples were HeLa cells stained with ABDS according to the protocol given in section 2.5. Unstained by ABDS HeLa cells were used as a control. Before the study, the cells were stained with DAPI dye, which penetrates only into the nuclei of dead cells. ^25^ Cell survival was analyzed using flow cytometry 2, 24, and 48 hours after staining. The results of the study are shown in Fig. 8. As can be seen in the figure, ABDS is not cytotoxic since the number of dead cells in stained samples is statistically indistinguishable from the number of dead cells in control samples.

**Figure 8:**
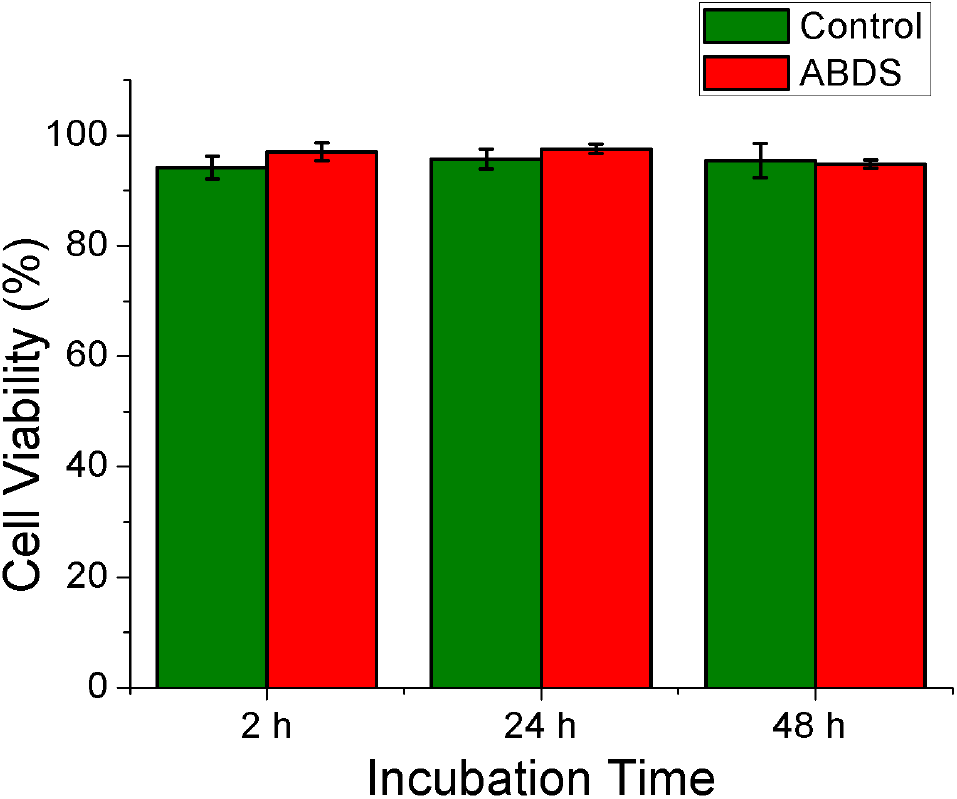
Cytotoxicity test using cell flow cytometry. The number of surviving cells in the control sample and stained by ABDS sample is statistically indistinguishable, therefore, ABDS is not toxic to cells. Error bars are calculated as 95% confidence intervals.

### 4.2 Exposure dose test

Various experiments with living cells, in particular in medicine and pharmacology areas, may require a long period of observation, including several days (a cell proliferation study, ^26^ *in vitro* drug testing, ^27,28^ *etc*.), which requires the dye to remain stable throughout the observation period. In this context, one of the most interesting properties of ABDS is its ability to not fade for a long time and even increase fluoresce brightness for several minutes under constant laser pumping. The problem of fading dye is often a significant obstacle to recording experiments in time-lapse mode and z-stack scanning of cells. In both cases, the initial frames are clear and contrast, but over time the typical dyes begin to fade, and, if the study requires a large number of frames, near the end of the experiment, the photographs become more blurry and significantly less informative and representative. In contrast to typical dyes, ABDS is very stable for long pumping times, as shown by the results of the exposure stress test, which we provide for ABDS and the commercial intravital dye DiBAC. During this test, we compared the ability of ABDS and DiBAC to maintain fluorescence at an acceptable level over long-term experiments. The stress exposure test consisted of continuously irradiating cells stained by ABDS and cells stained by DiBAC with a Zeiss fluorescence microscopy pumping laser (see Sec. 2.6) with emission wavelength 488 nm. This wavelength excites the DiBAC dye, and we also used it to pump ABDS to ensure that the dyes were on equal terms. The cells were continuously illuminated by the laser emission for 20 min except time periods when photograph occurred. The photographs were collected using 1 sec exposure.

The obtained results are presented in Fig. 9. The photographs were processed using the settings “Best-Fit” of the program AxioVision. As can be seen from photographs sequences, over time about 10 minutes after exposure the fluorescence of DiBAC becomes less bright and, as a result, the pictures become less clear.

**Figure 9:**
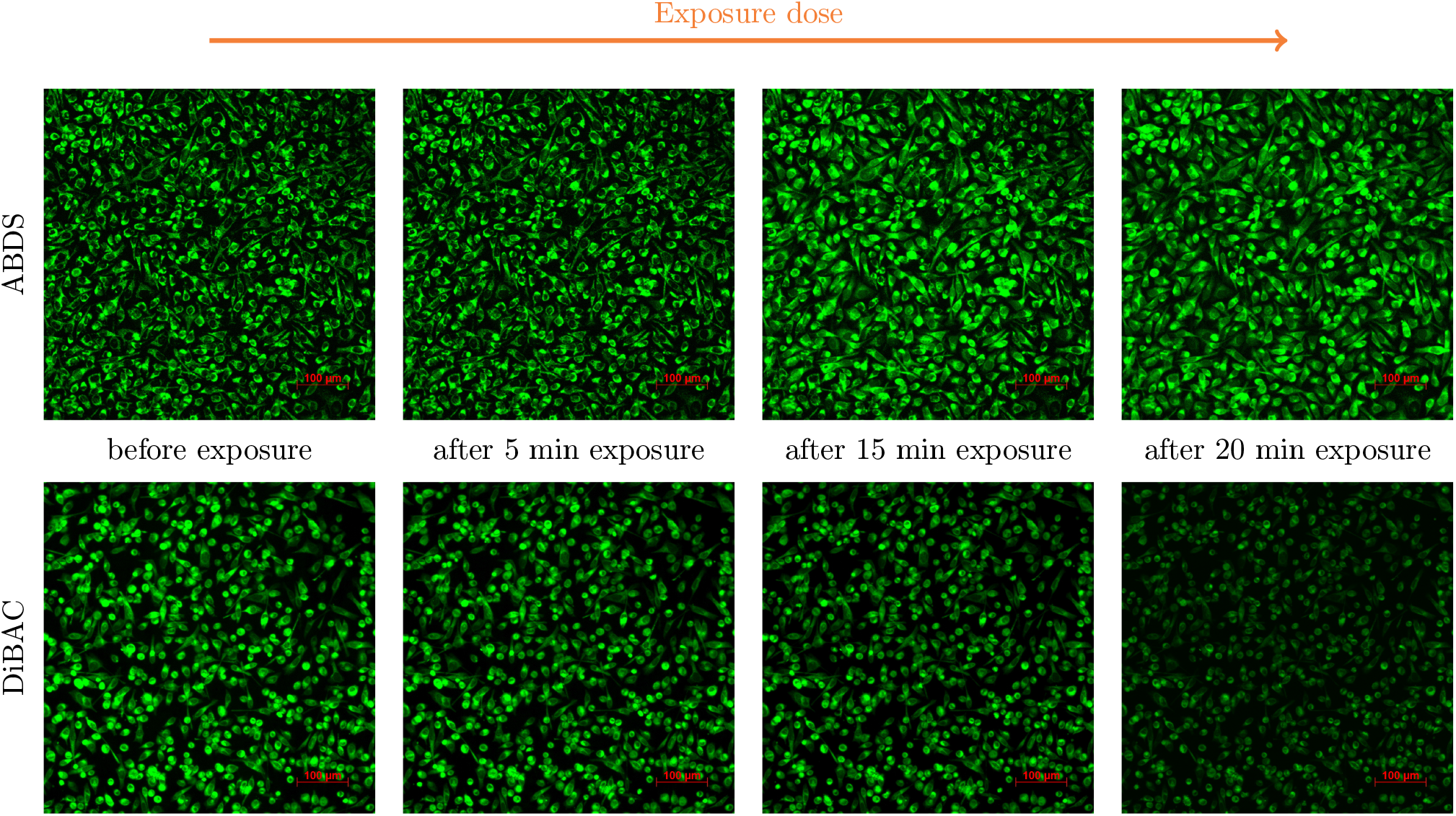
Exposure dose stress test for ABDS and DiBAC. It can be seen that after 15 minutes the fluorescence of the ABDS dye becomes brighter and then after 20 minutes it fades slightly. In contrast, fluorescence of the DiBAC dye just becomes less brighter with the time and, as a result, the photograph becomes less clear.

This problem with fluorescence becomes especially significant when careful comparison of samples before and after the test treatment is necessary, since it is impossible to tell whether the changes are due to the experimental treatment or to dye burnout.

In contrast, the dependence of the fluorescence of the ABDS on the exposure dose is not trivial. ABDS fluorescence initially increases for about 15 minutes and then the fluorescence becomes less bright. However, it should be noted that after 20 minutes of the experiment, the fluorescence did not drop below the initial level. This useful property of the ABDS dye can be applied for many applications, for example to observe cell division, apoptosis process.

The nature of the increasing ABDS intensity can be explained by quantum quenching phenomena. ^29^ Indeed, when the concentration of ABDS is high, then the emitted photons from one center have the high possibility to absorb in another, where it can relax in a nonradiative way. However, during the pumping process some fluorescence centers start to burn down, which decreases the overall dye concentration and thus decreases the probability of the nonradiative relaxation. The latter led to an increase of the total ABDS fluorescence intensity.

### 4.3 Selective staining

By conducting experiments with ABDS, we found that this dye can stain different types of cells in different ways. In particular, we observed that ABDS stains HeLa cells brighter than RIN m5F – rat insulinoma insulin-producing cells. ^30^ This statement is supported by the data presented in Fig. 10. In this experiment, we incubated HeLa cells with RIN m5F in the same Petri dish, after which we applied ABDS to such a heterogeneous cell culture. From panel (a), which represents the fluorescence signal in pseudocolor, it can be seen quite clearly that the fluorescence intensity is higher for the fusiform-shaped cells, which we attribute to HeLa, in contrast to RIN m5F cells, which are mainly spherical. Panel (b), which is a combination of the light-transfer mode image with the ABDS signal added to the green channel, also confirms the selective staining of ABDS. Thus, ABDS has the potential to create protocols for distinguishing different cells from heterogeneous cultures.

**Figure 10:**
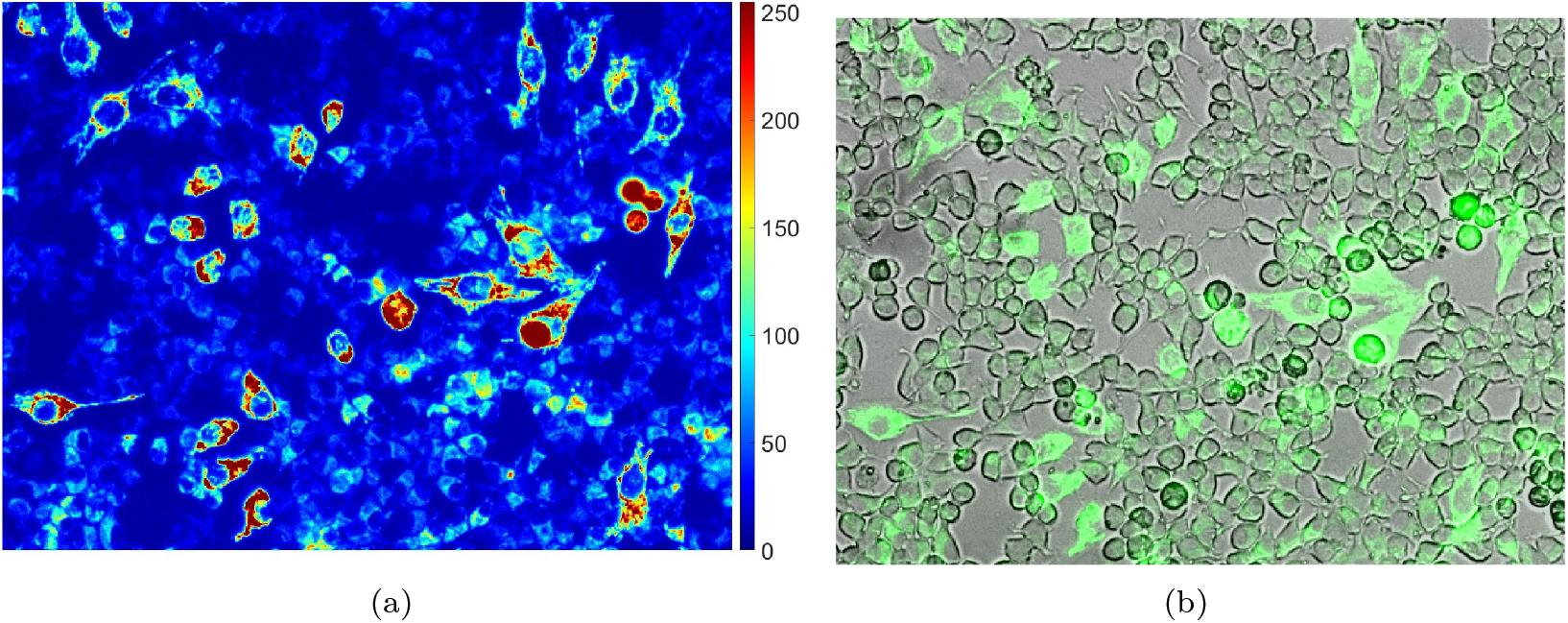
Demonstration of selective ABDS staining for a test heterogeneous HeLa/RIN m5F culture. (a) ABDS fluorescence signal presented in pseudocolor, (b) transfer light (bright field) mode image with ABDS fluorescence superimposed on the green channel. The obtained data shows that fusiform-shaped HeLa cells have a higher ABDS signal than spherical RIN m5F, indicating that ABDS can be used to distinguish between different cell lines.

### 4.4 High Detalization

Another advantage of ABDS is its ability to stain focal contacts – the protein processes of adhesion cells, which they use to attach to the surface of the culture dish. ^31^

Of the commercially available dyes, the Deep Red Cell Mask has the ability to provide a sufficiently detailed picture. A comparison of photographs obtained using a Zeiss Observer.Z1 confocal microscope for Deep Red Cell Mask and ABDS dyes is shown in Fig. 11. As can be seen in these photographs, ABDS paints focal contacts as well as Deep Red Cell Mask, despite the price difference of more than a hundred times (see Tab. 1).

**Table 1:** Comparison of the cost of various intravital cell membrane dyes.

| Dye | Cost for 100 μL |
| --- | --- |
| ABDS | <\$2 |
| DiBAC | \$40 |
| Deep Red | \$320 |

**Figure 11:**
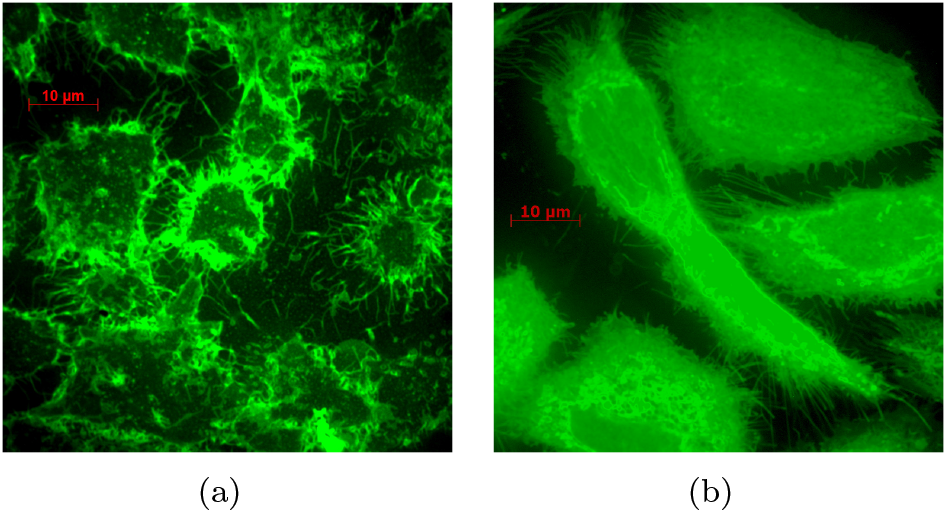
The lowest lay of Z-stack image shows the structure of focal contacts: (a) – Deep Red Cell Mask; (b) – ABDS. The presented data again justify the sub-micron resolution of the ABDS (cmp. with Fig. 5).

Another interesting property of ABDS was discovered when it stained RIN m5F cells. It was found that RIN m5F cells are unevenly stained, some cells’ fluorescence are brighter and some are paler.

Although in this study we did not determine to which part of the cell membrane ABDS binds, but from Fig. 12 it is evident that when staining RIN m5f, the dye binding targets are distributed unevenly, in contrast to HeLa cells. This may be due to the different number of ABDS binding targets depending on the phase of the cell cycle.

**Figure 12:**
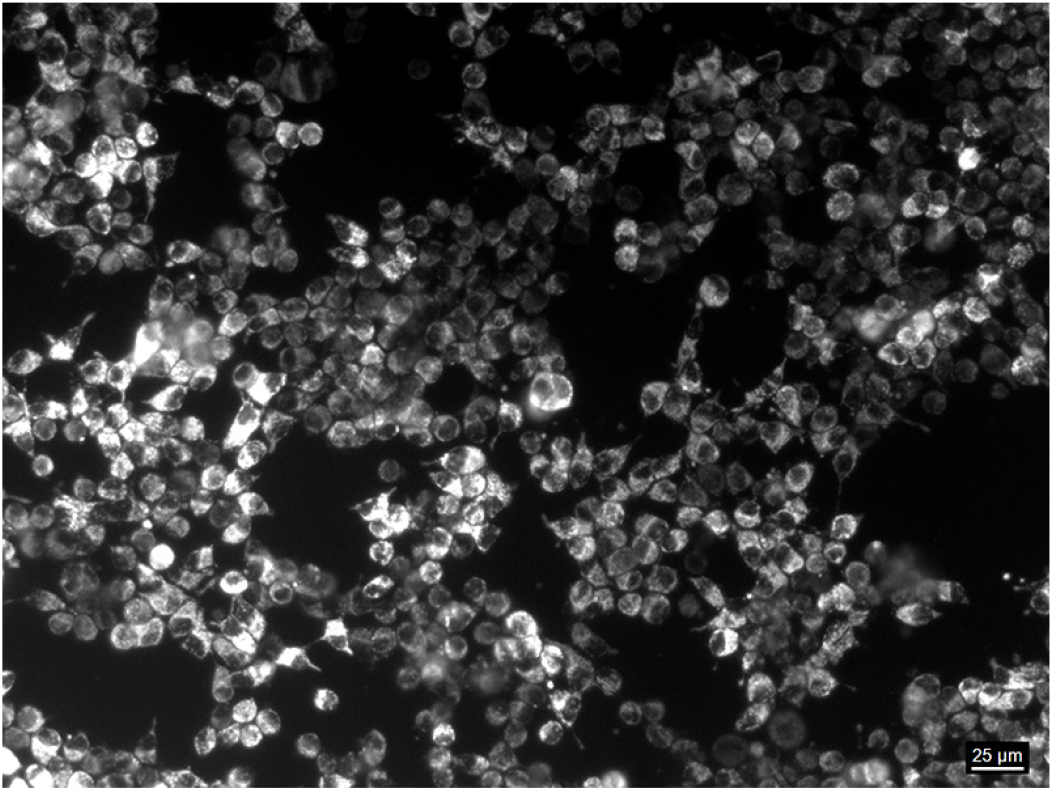
RIN m5f cells dyed by ABDS. The fluorescence of cells is heterogeneous, which may indicate their different states.

### 4.5 ABDS cost-effectivity

As mentioned earlier, one of the most important features of ABDS is its low-cost. Unlike expensive commercial cell dyes, ABDS can be manufactured in any laboratory and even in the field environment because the ingredients necessary for its manufacture – red permanent marker, ethanol, and PBS, are absolutely affordable.

The permanent marker Edding 140S costs less than $2 and one pen can be used to prepare at least 100 ml of ABDS. At the same time, the cost of commercial dyes is significantly higher. For example, DiBAC, also used for intravital staining of cell membranes, costs about $40, and the cost of another intravital dye, Deep Red Cell Mask, reaches $320, as can be seen in Tab. 1.

In order to obtain the same amount of ABDS as other dyes listed in Tab. 1, it is necessary to dilute the approximately 100 mm line drawn with the marker according to the protocol described in Sec. 2.1. However, to prepare the ABDS, one permanent red marker and ethanol must be purchased for dilution, so the cost of preparing the dye can be estimated at $2.

## 5 Applications for cell research

As can be seen in the previous Sec. 4.2, the properties of ABDS allow it to be used for experiments that require long-term observation or a high frame rate. In this section, we demonstrate several studies showing the benefits of ABDS applications.

### 5.1 Z-stack and high resolution of ABDS

The property of ABDS not to lose its fluorescence over time can also be used to obtain a three-dimensional image of a cell at high magnification. When studying cells with magnification 68X, the effect of fluorescence enhancement occurs faster than lower magnification, within a few seconds, so it does not have a significant effect on the quality of the resulting Z-scan of the cell. ABDS staining allows one to obtain highly detailed images of cells, for example, in Fig. 13 cell morphology, including focal contacts, is clearly visible.

**Figure 13:**
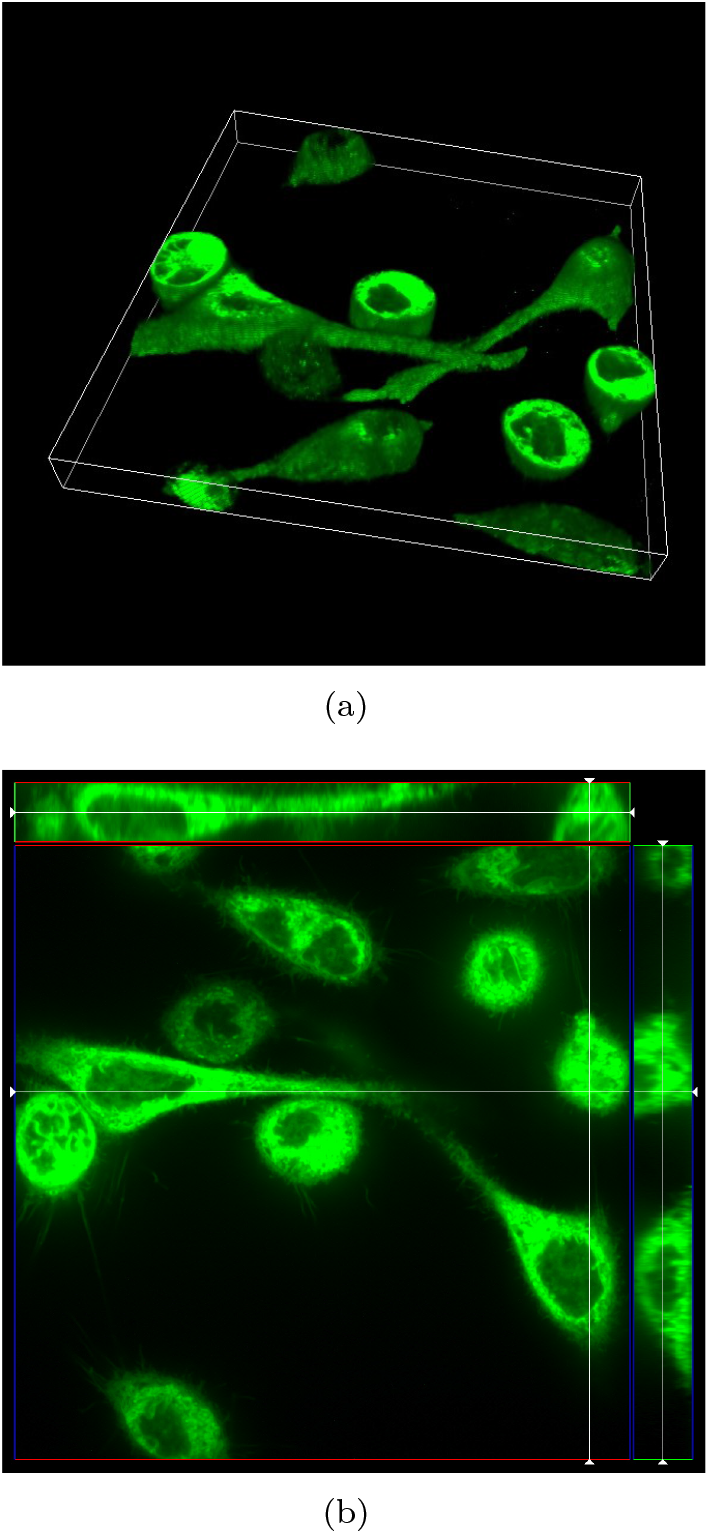
HeLa cells Z-stack 3D tomography (a) and cut-view (b) obtained with ABDS. One can note the absence of dye fading.

### 5.2 Prolifiration study

To illustrate the benefits of ABDS, we conducted the following experiment. HeLa cells were stained with ABDS and scattered on fluorescent squares made of photoresist AQUA MER ME720 (Zhuhai Dynamic Technology, China) for easy visualization of their growth (Fig. 14). Sodium butyrate, known to have cytostatic properties, ^32,33^ was added to the experimental samples. No sodium butyrate was added to the control samples. The purpose of this experiment was to demonstrate that sodium butyrate would inhibit cell growth, ^34,35^ while in control samples, cell growth should follow the cell growth curve.

**Figure 14:**
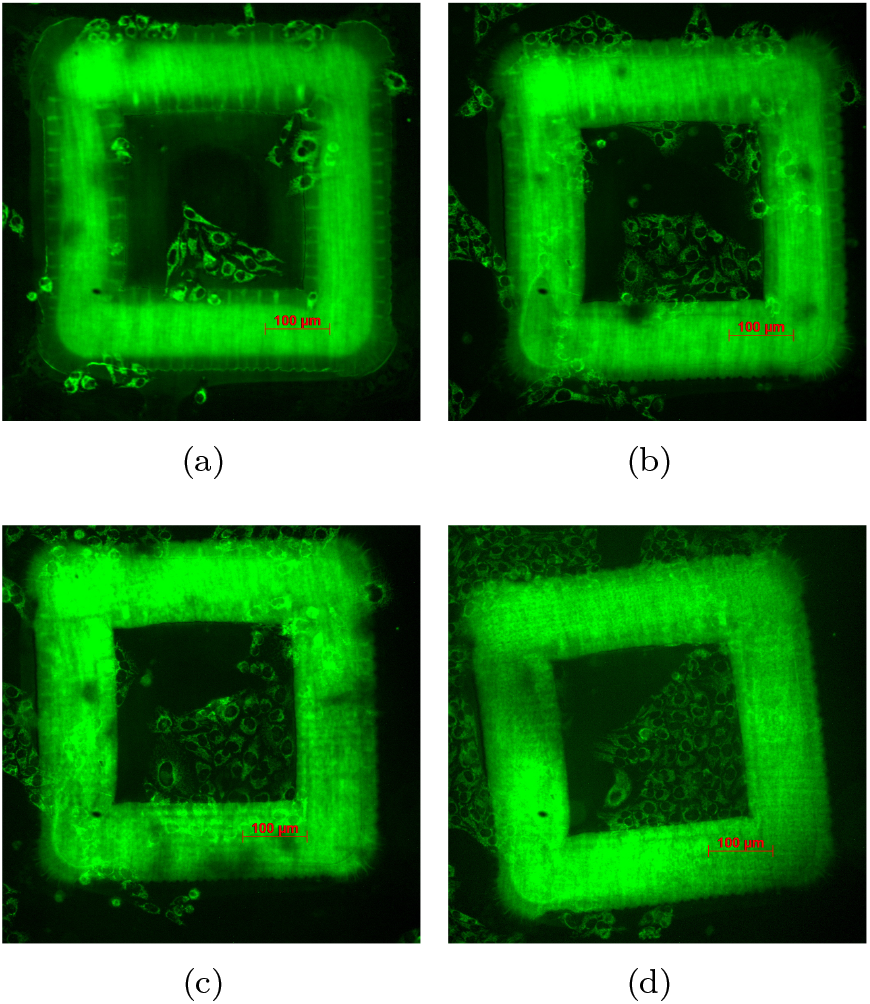
Proliferation study of HeLa cells stained by ABDS. (a) – 1 day after staining, (b) – 2 days, (c) – 3 days, (d) – 4 days. One can see that during cells division process ABDS passes to daughter cells membrane.

As can be seen from the graph presented in Fig. 15 the number of HeLa cells incubated with sodium butyrate decreased rather than remained constant as expected in this experiment, which may be due to the natural process of cell death. At the same time, HeLa cells, to which cytostatic was not added, actively divided. Thus, we have shown that ABDS can be used to study cells proliferation activity. Furthermore, the data obtained justify not only that ABDS is a non-toxic intravital dye for cells (compare with Sec. 8), but also that it does not have a significant effect on the process of their division.

**Figure 15:**
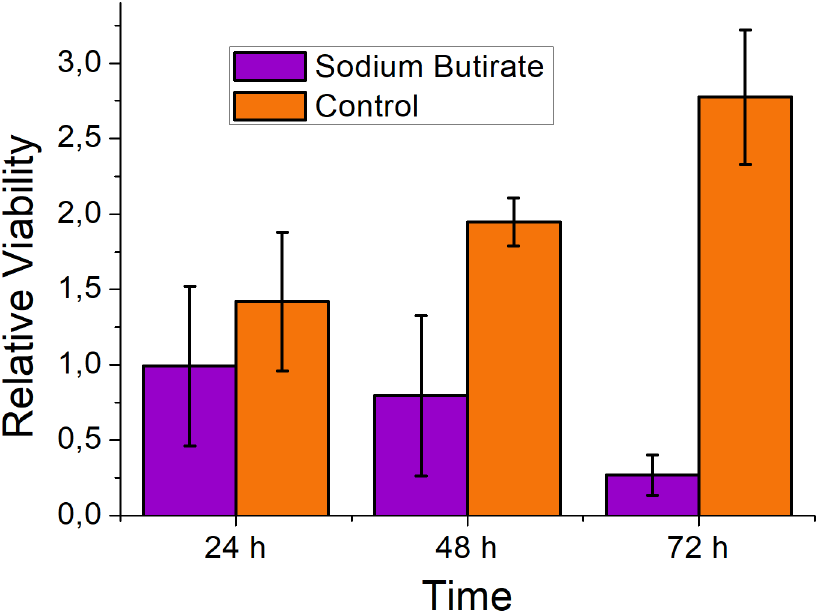
Proliferation study results: during the experiment the number of the cells in counting squares (Fig. 14) has grown for control cells group, and has decreased for cells treated with sodium butyrate.

### 5.3 Adhesion study

As an example of using ABDS for rapidly occurring phenomena in the study of cells, we consider an experiment with the de-adhesion of HeLa cells under the action of a trypsin-Versene solution (Biolot, Russia). The trypsin-Versene solution acts on the so-called focal contacts that attach the cell to the surface of the culture dish. As a result of exposure, the cell changes its fusiform shape to a spherical one. ^36^

The results of the study of Hela cell de-adhesion under the action of the trypsin-Versene solution are shown in Fig. 16. It can be clearly seen that, at the initial time before the addition of the trypsin-Versene solution, cells were attached to the surface using focal contacts and have a fusiform shape. Furthermore, after a solution of trypsin-Versene was added, the focal contacts were destroyed, and the cell acquired a spherical shape. As can be seen in Fig. 16, during such an experiment the ABDS does not reduce its fluorescence, which makes it possible to study in detail the process of cell de-adhesion.

**Figure 16:**
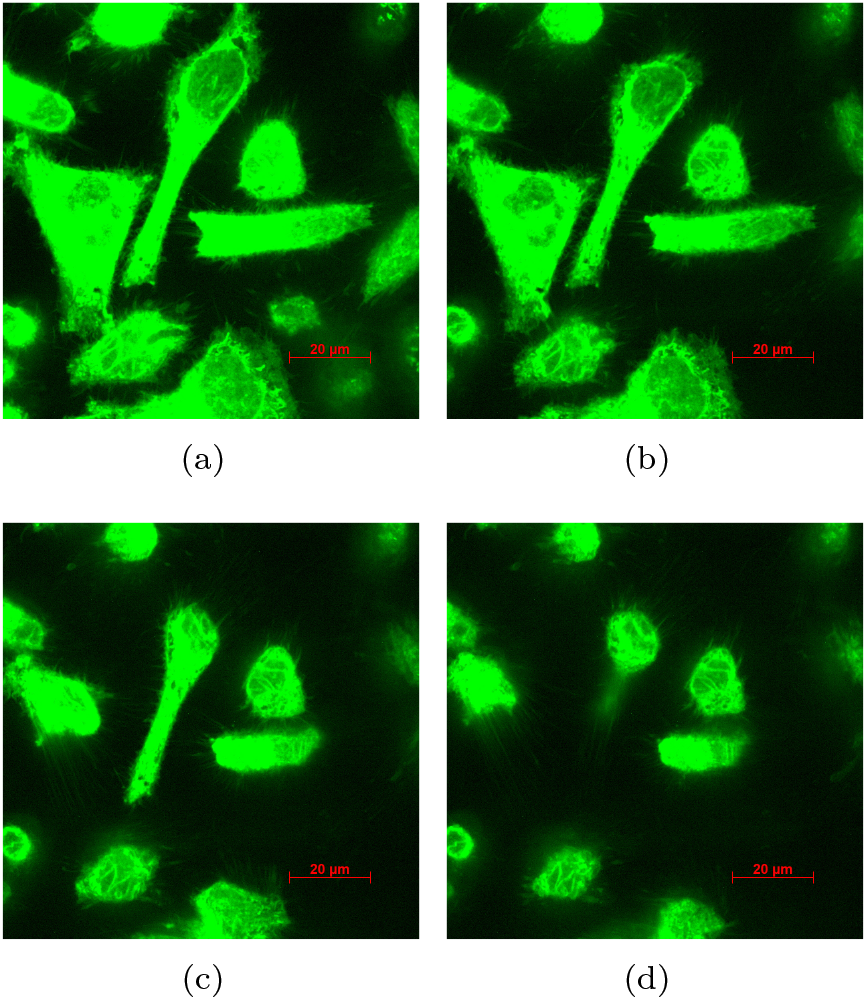
De-adhesion of HeLa cells after trypsin-Versene solution added. (a) – Before trypsin-Versene addition, (b) – 30 s after addition, (c) – 1 min 30 s after addition, (d) – 2 min 30 s after addition. One can see, the ABDS allows to monitor the cells form in real-time regime.

### 5.4 Use of ABDS in impedance spectroscopy

One of the fastest growing areas in bioelectronics is the development of impedance cytosensors. ^18,37–40^ Such devices convert a biological signal received directly from cells into an electrical signal, which allows study of a wide range of cell phenomena. ^41^ Typically, the impedance cytosensor is a modified Petri dish with an array of microelectrodes on the bottom – multielectrode array (MEA). ^42,43^ Since the impedance of the electrode on which living cells are located depends significantly on their state, ^44–46^ for the development and optimization of impedance cytosensors external monitoring of cells is required. Because most of the materials used to make MEA electrodes are opaque, ^47^ the convenient method to solve this problem is to use fluorescence microscopy methods and various dyes to visualize cells. The dye required for such a study must meet both biological and physical criteria: firstly, it must be intravital and the cells must not change their state after its addition, and, secondly, such a dye must not stain the surface of the MEA. As was shown in Sec. 4.1, ABDS does not affect cell viability and shape; therefore, ABDS is a promising candidate for a low-cost and reliable dye for bioelectronic devices. To justify this statement, we have conducted an experiment for the cell de-adhesion study using MEA and impedance spectroscopy (cmp. with Sec. 5.3).

In Fig. 17 the results of the study of de-adhesion of a cell population with the 60StimMEA200/30-Ti multielectrode array (Multichannel Systems, Germany) are presented, and in Fig. 18 the results of the study of de-adhesion of a single cell with usage of the 60MEA200/30R-Ti multielectrode array (Multichannel Systems, Germany) are shown. The impedance data were collected using the setup presented in Refs. ^18^ and. ^48^ Control electrodes in the photographs are high-lighted in red, and cell-covered electrodes are high-lighted in green.

**Figure 17:**
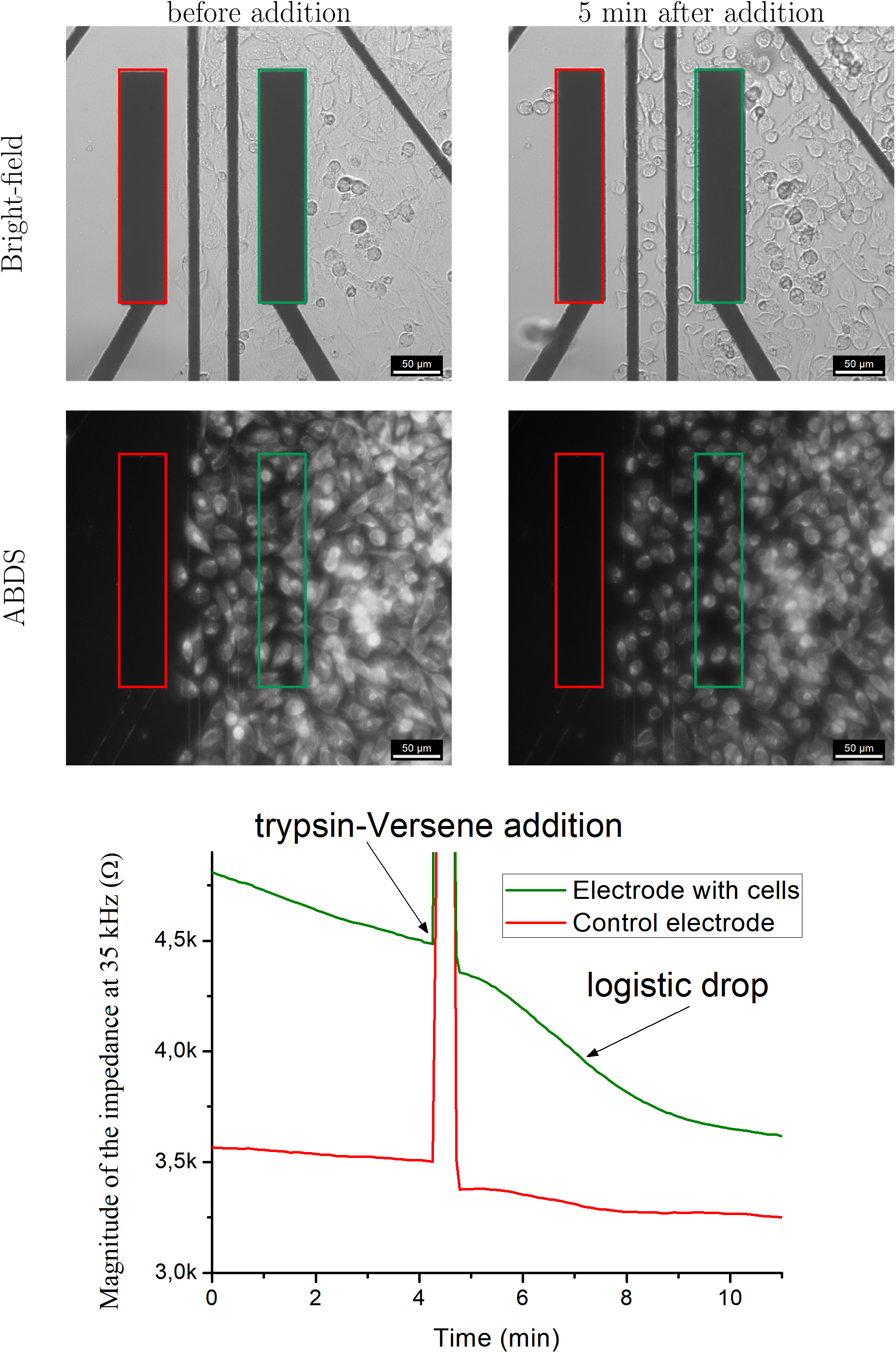
Cell population de-adhesion study by impedance spectroscopy using ABDS dye. It can be seen that while the cells change their shape from fusiform to spherical under the action of trypsin-Versene, the impedance of the cell-coated electrode decreases in accordance with the with Giæver-Keese model. ^18,45^

**Figure 18:**
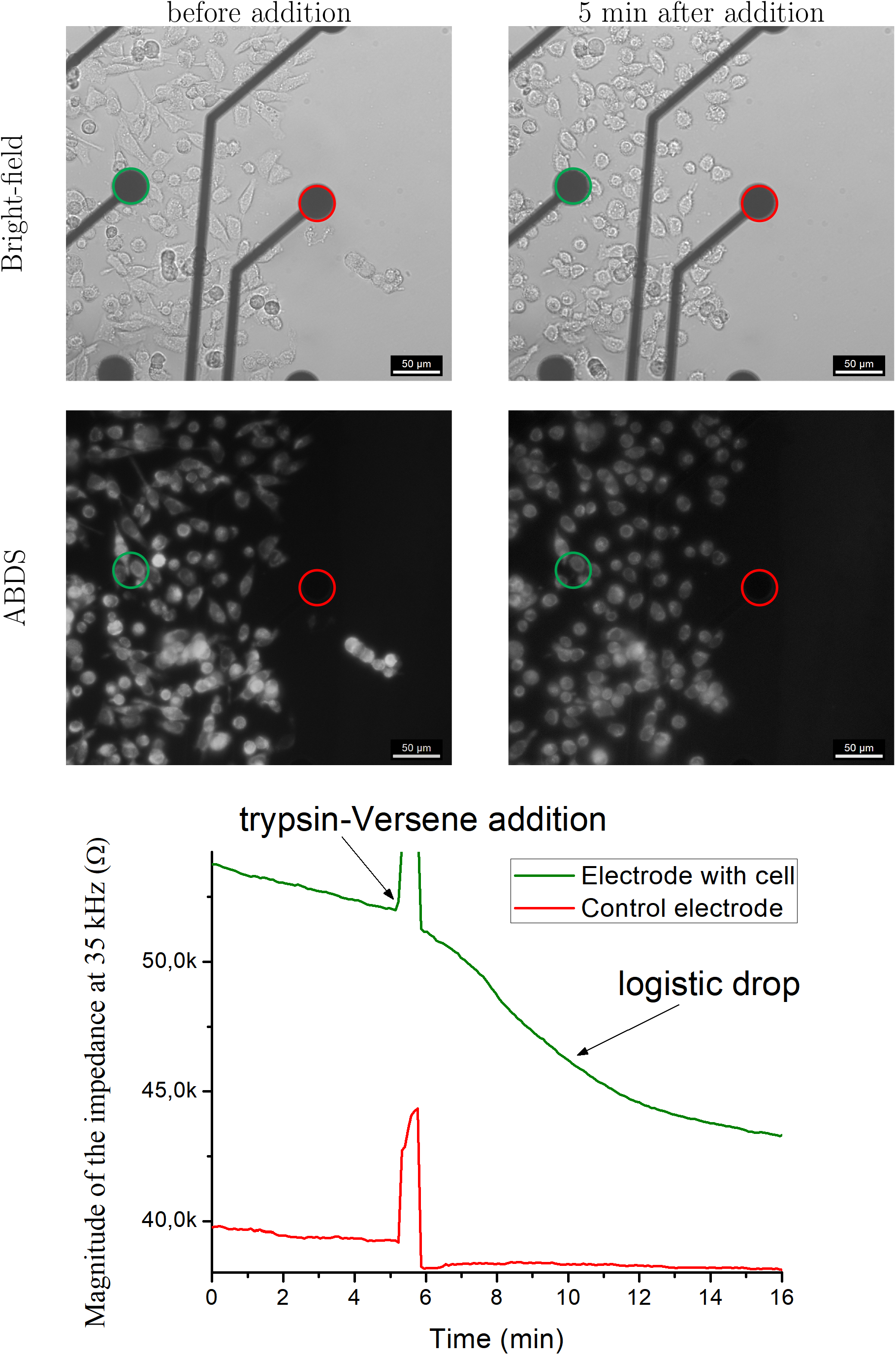
Single cell de-adhesion study by impedance spectroscopy using ABDS dye. The impedance evolution of the the coated by cells electrode and dynamics of the cells shape are similar to the case of the large electrodes (Fig. 17).

As can be seen from the presented data, ABDS allows clear visualization of cells located on electrodes, both in the case of studying cell populations and in the case of studying single cells. In addition, it is clearly seen that the shapes of the cells before and after the experiment are significantly different, and this phenomenon explains the shape of the curve in the results of the cell impedance study. The cells acquire a spherical shape, and the area of contact of the cell with the electrode decreases, therefore, the impedance of the electrode with the cells also decreases in both cases. We also observed that ABDS does not change the MEA electrodes’ impedance level significantly, and it does not stain the MEA dish. Thus, we have shown that ABDS can be used in bioelectronic devices to visualize cells on the surface of electrodes, which is important for the development of new biosensors.

## 6 Conclusion

In this study, we have developed the DIY low-cost cell membrane dye ABDS, which is based on the permanent marker inc. We have investigated its properties such as biocompatability, emission, absorption, and Raman spectra. Based on the data obtained, we can conclude that ABDS is non-toxic and contains Rhodamine 6G fluorophore. The data presented in Sec. 4.4 and Sec. 5 demonstrates that ABDS can be used to obtain detailed cell images, which allows to study cell adhesion, proliferation activity, and bioelectrical properties. Separately, we note that the problem of different coloring by inks from different manufacturers mentioned in ^13^ does not arise in our case, and ABDS made from any permanent red markers colors the cells equally. Although only a few examples of the use of ABDS are presented in this paper, they show that ABDS is a promising candidate for the role of a cheap and accessible dye that can be easily used in the field because it does not require either careful sample preparation or special storage conditions. We hope that the method of intravital cell staining with ABDS proposed in this work will significantly accelerate the development and testing of new bioelectronic devices.

## 7 Acknowledgments

The authors express their gratitude to Metelkina E. M., Ph. M. Dubina, Filatov N. A., Blinova M. I., and Dubina M. V. for comprehensive assistance and support. This study was carried out with the support of the Ministry of Science and Higher Education (Project № FSRM-2024-0001).

## Notes

### Competing Interest Statement

The authors have declared no competing interest.

